# Dose-dependent, opioid-induced sleep disruption is partially mediated by motor activity but not by body temperature in C57BL/6J mice

**DOI:** 10.1101/2025.06.22.660933

**Authors:** Diana Zebadúa Unzaga, Natania W. Cooper, Charlie E. Olson, Xiaojuan Zhu, Michael A. O’Neil, Joshua M. Price, Ralph Lydic, Helen A. Baghdoyan

## Abstract

States of sleep and wakefulness, motor activity, and body temperature are temporally linked, yet these variables are often studied independently. Here we report novel results from male C57BL/6J mice (n = 24) that quantify the causal and correlational relationships between these temporally linked measures. The initial experiments used 12 mice to identify the lowest antinociceptive doses of fentanyl (0.1 mg/kg) and morphine (1 mg/kg). Twelve additional mice were implanted with telemeters to simultaneously record EEG, electromyogram, motor activity, and subcutaneous body temperature. Doses of fentanyl (0.1, 0.3, 1, 3 mg/kg) and morphine (1, 3, 10, 30 mg/kg) caused significant (P<0.05) increases in wakefulness and decreases in NREM and REM sleep. Fentanyl (0.1 to 3 mg/kg) and morphine (3 to 30 mg/kg) significantly increased motor activity. Body temperature during wakefulness was significantly decreased by fentanyl (1, 3 mg/kg). Morphine increased (3 mg/kg) and decreased (30 mg/kg) body temperature. Mediation analyses showed that the increase in wakefulness caused by fentanyl and morphine was partially mediated by motor activity, but not by changes in body temperature. These results provide the first complete dose-response data for effects of fentanyl and morphine on simultaneously acquired measures of sleep/wake states, body temperature, and motor activity. Compared to human data, these results from mice reveal similarities (sleep disruption, hyperthermia and hypothermia) and differences (increased motor activity) caused by fentanyl and morphine. The results also demonstrate the power of mediation analyses for providing nuanced insights into opioid effects on clinically relevant neurobehavioral phenotypes.

**Significance Statement:** Sleep disruption is an undesired, clinically significant effect of opioids prescribed to manage pain. Sleep disruption exacerbates pain and can lead to increased opioid requirement, as well as increased risk of addiction relapse. The significance of this study is the discovery that in mice, even the lowest doses of fentanyl and morphine that caused antinociception also caused sleep disruption. Thus, in mice, it is not possible to provide antinociception using fentanyl or morphine without causing sleep disruption.

## Introduction

Despite the ongoing crisis of addiction, opioids remain a vital tool for managing acute (Dowell et al., 2022; Collins et al., 2025) and chronic (Dahlhamer et al., 2018; Glare et al., 2025) pain. Clinical administration of opioids for pain management is complicated by opioids causing respiratory depression (Ramirez et al., 2021; Eckert and Yaggi, 2022; Baldo, 2025; Cipriano et al., 2025) and having abuse potential (Duerksen et al., 2025; Marchette et al., 2025; Volkow and Dye, 2025). The present study was designed to focus on clinically relevant phenotypes produced by anatomically distributed neuronal networks. Accordingly, one goal of this study was to address a gap in knowledge by quantifying the dose-dependent effects of acutely administered fentanyl and morphine on sleep/wake states and state-dependent motor activity and body temperature. This study also sought to determine whether there are doses of fentanyl and morphine that produce the desired effect of antinociception without causing the undesired effect of sleep disruption. A systematic review of the literature found no previous, complete dose-response data comparing the effects of fentanyl and morphine on nociception, motor activity, and sleep/wake states in C57BL/6J (B6) mice.

Sleep disruption is a clinically significant, unwanted effect of fentanyl and morphine (Eacret et al., 2020; Lydic et al., 2022). Among humans, reduced sleep causes neurobehavioral deficits (Yamazaki et al., 2021) and increases the risk of addiction relapse (Brower and Perron, 2010; Roehrs et al., 2021; Eckert and Yaggi, 2022; López-Muciño et al., 2022). Sleep deficiency is comorbid with many psychiatric disorders and insomnia may be a risk factor for developing depression (Atwood, 2022). Sleep disruption delays wound healing in mice (McLain et al., 2018) and humans (Anderson and Jeter, 2023). Sleep loss causes hyperalgesia in humans (Roehrs et al., 2006; Finan et al., 2013), rats (Muncey et al., 2010; Vanini et al., 2014), and mice (Dalanon et al., 2021). Hyperalgesia increases analgesic requirement (Fletcher and Martinez, 2014; Knutson, 2015), creating a vicious cycle for pain management (Haack et al., 2020).

The temporal organization of wakefulness, rapid eye movement (REM) sleep and non-REM (NREM) sleep is essential for sleep, just as the normal cycle of inspiration and expiration is required for gas exchange. The disruption of human sleep by intruding periods of wakefulness is common for first responders, medical residents, and caregivers of neonates. Among adults, sleep fragmentation disrupts sleep continuity and can result in severe health consequences (Thomas, 2024). Human sleep fragmentation is associated with sleep disordered breathing and daytime hypertension (Morrell et al., 2000). In humans (Djonlagic et al., 2021) and mice (Li et al., 2022) sleep fragmentation causes cognitive impairment. Opioids also cause dose-dependent sleep fragmentation in humans (Eacret et al., 2020; Greenwald et al., 2021; Lydic et al., 2022; Akinnusi et al., 2024).

The foregoing findings raise the question of whether it is possible to achieve opioid-induced antinociception without also causing significant sleep disruption. To test this hypothesis, we obtained simultaneous measures of sleep/wake states, motor activity, and subcutaneous body temperature from B6 mice before and after administering eight doses of fentanyl and eight doses of morphine. The choice of dependent measures was informed by previous studies showing that in mice opioids increase locomotor activity (Mickley et al., 1990; Varshneya et al., 2019; Haouzi et al., 2021; Santos et al., 2022) and disrupt sleep (O’Brien et al., 2021; Gamble et al., 2022; Sharma et al., 2024). Opioids also alter body temperature in mice (Rosow et al., 1980; Baker and Meert, 2002) and humans (Little and Kosten, 2023).

Fentanyl and morphine produce dose-dependent changes in neurobehavioral and autonomic functions that reflect actions on opioid receptors throughout the central and peripheral nervous system. Computational approaches of systems pharmacology, including mediation analyses, are increasingly used to quantify and reveal direct and/or indirect drug effects arising from anatomically distributed but interacting neuronal networks (Bai and Abenethy, 2013; Yamamoto et al., 2015; Whibley et al., 2019). Mediation analysis, a specific application of structural equation modeling, is a statistical approach used to quantify the extent to which an independent variable (predictor) influences a dependent variable (outcome) via one or more intervening mediator variables. In the present study mediation analysis was used to quantify the extent to which opioid-induced sleep disruption was mediated by changes in motor activity and/or body temperature. These analyses revealed that increased wakefulness caused by fentanyl and morphine was partially mediated by motor activity but not by changes in body temperature.

## Materials and Methods

### Animals

All experiments involving animals were reviewed and approved by the University of Tennessee Institutional Animal Care and Use Committee and were conducted in accordance with the ARRIVE guidelines (Percie du Sert et al., 2020). Adult, male B6 mice (Stock# 000664; n=24) were purchased from The Jackson Laboratory (Bar Harbor, ME). At the University of Tennessee Knoxville mice were housed on a 12 h:12 h light-dark schedule. Temperature (20.6 to 24.5 °C) and humidity (30 to 70%) were maintained during 10 to 12 air changes every hour. Mice had *ad libitum* access to water and rodent chow (Teklad 8640, Envigo, Madison, WI). Igloos, nestlets, and paper cones were rotated in home cages to provide environmental enrichment. Before surgical implantation of telemeters, littermates were housed together with no more than five mice in each cage (19 x 29 x 13 cm). Cages, enrichment items, and bedding (Harlan Soft Cob Enrichment Bedding) were changed once per week. Mice were monitored daily by laboratory members and by sensors that provided the investigators with 24/7 wireless receipt of temperature and humidity data. Mice were given at least one week to adjust to the housing environment before starting nociceptive testing or surgical procedures.

### Quantifying antinociceptive effects of fentanyl and morphine using the tail withdrawal latency test

These experiments were designed to quantify the range of antinociceptive opioid doses to be used for subsequent studies of sleep. One week prior to measuring tail withdrawal latency, mice were conditioned daily to being handled and placed in an acrylic restraint tube with their tails extending outside of the tube. For data collection, the distal third of the tail was placed in a warm water bath (47.8 to 50°C). Nociception was measured as the time (s) from tail immersion into the warm water to the time of mouse-initiated tail removal from the warm water (Gårdmark et al., 1998; Schildhaus et al., 2014). To prevent tissue damage, the maximum amount of time that mouse tails remained in the warm water was 15 s, after which the experimenter removed the mouse tails from the water bath. Thus, the maximum possible value for tail withdrawal latency was 15 s (Glovak et al., 2017). Nociceptive testing began by obtaining baseline (no injection) measures of tail withdrawal latency for each mouse. Mice then received subcutaneous injections (0.3 mL) of saline (vehicle), fentanyl, and morphine. The mean ± SD age of the mice (n=12) used for nociceptive testing was 26 ± 5.2 weeks and mean ± SD body weight was 35 ± 4.9 g.

### Chronically implanted telemeters enabled simultaneous recording of EEG, electromyogram (EMG), motor activity, and subcutaneous body temperature

Twelve additional B6 mice naïve to opioids were implanted with telemeters (HD-X02, Data Sciences International (DSI), New Brighton, MN). Telemeters were used to quantify effects of opioids on states of sleep and wakefulness, motor activity, and body temperature from freely behaving mice in their home cages. Surgeries were performed under general anesthesia with 2% isoflurane (Henry Schein, Melville, NY) delivered in 100% oxygen at a flow rate of 1 L/min. At the time of surgery mean ± SD mouse age was 22 ± 3.2 weeks and mean ± SD mouse body weight was 33 ± 4.2 g. Isoflurane delivery via an agent-specific vaporizer was monitored by spectrophotometry (Cardiocap/5; Datex-Ohmeda Inc., Madison WI).

Mice were placed in a plexiglass induction chamber and loss of righting reflex was used as an index of loss of consciousness (Vanini et al., 2014). Mice were shaved over the skull and cranial half of the dorsum. Anesthetic depth was assessed based on breathing rate and lack of response to a foot pad pinch. Mice were then placed in a stereotaxic frame (David Kopf Instruments, Tujunga CA; model 962) fitted with a mouse adaptor (model 921) and a mouse anesthesia mask (model 907). Ketoprofen (5 mg/kg) in saline (0.3 mL) was administered subcutaneously to provide post-operative analgesia and prevent dehydration. A subcutaneous radio-frequency identification chip (ID-100B 1.4, MICROCHIPID, Lake Zurich, IL) was implanted to provide unique alphanumeric identification for each mouse. Throughout surgery the eyes of the mouse were protected with ophthalmic ointment. Core body temperature (36–37 °C) was monitored via a rectal thermometer and maintained using hot water circulating through a leakproof pad (Gaymar TP400 T/Pump heat therapy system). The skin was prepared aseptically for surgery by alternating iodine solution and iodine scrub. Respiratory rate and body temperature were measured every 10 min until the end of the surgical procedure.

A midline skin incision was made beginning at the top of the skull and extending caudally between the scapulae. Bipolar EEG recording electrodes originating from the telemeter were connected to stainless steel screws (Antrin Miniature Specialties, Fallbrook, CA) implanted above the cortex. Burr holes for the screws were made at stereotaxic coordinates 1.0 mm anterior to Bregma and 1.0 mm lateral to the midline (negative lead), and 3.0 mm posterior to Bregma and 3.0 mm lateral to the midline (positive lead) (Paxinos and Franklin, 2019). Two electrodes were placed in the cervical trapezius muscle in the dorsal region of the neck to record the EMG. The telemeter was placed in a deep subcutaneous pocket in the right dorsal flank. The dorsal portion of the incision over the skull and neck was closed with non-absorbable sutures (Prolene 6-0, Henry Schein, Melville, NY) in a simple interrupted pattern. The caudal portion of the incision was closed with wound clips (Henry Schein). Mice recovered in a heated cage and were monitored continuously until they displayed normal righting and ambulation.

After surgery, mice were housed in individual cages to prevent disruption of the implant by cage mates. Daily health examinations were performed by investigators to assess mouse well-being. Postoperative monitoring focused on wound healing and behavioral indices of health including normal locomotion, nest building, and normal gain of body weight. Mice recovered during a two-week postsurgical healing period that preceded data collection.

### Experimental design

A within-subject, repeated measures design was used to reduce the number of animals needed (Piantadosi, 1997). Each mouse received subcutaneous injections (0.3 mL) of saline (vehicle control) and increasing half-log doses of fentanyl (0.001 to 3 mg/kg) and morphine (0.01 to 30 mg/kg). Injections into the same mouse were separated by three to four days. To reduce the potential for order bias, each mouse was randomly assigned to receive either fentanyl or morphine treatment first. After each mouse received saline and all doses of the first opioid, mice were given at least one week of no injections before starting the second opioid. A second saline injection was administered before the second treatment protocol was started. Results of the first and second saline treatment were compared to confirm that the dependent measures of nociception, sleep/wake states, motor activity, and body temperature had not changed due to treatment or time. Mice were weighed and given health examinations before each experiment.

### Recording and quantifying states of sleep and wakefulness, motor activity, and subcutaneous body temperature

Recordings of the dependent measures were obtained from mice in their home cages by placing each cage on a receiver that communicated with the data acquisition computer (DSI Matrix 2.0 MX2). Subcutaneous injections were given within 90 min of light onset and 4-h recordings began immediately after each injection. EEG and EMG signals were used to objectively quantify states of wakefulness, NREM sleep, and REM sleep. Motor activity was measured by the telemeter as activity counts in arbitrary units (a.u.). The implanted telemeter also measured subcutaneous body temperature in degrees centigrade (°C). EEG, EMG, motor activity, and body temperature were recorded simultaneously (Fig. 1A).

**Fig. 1.**
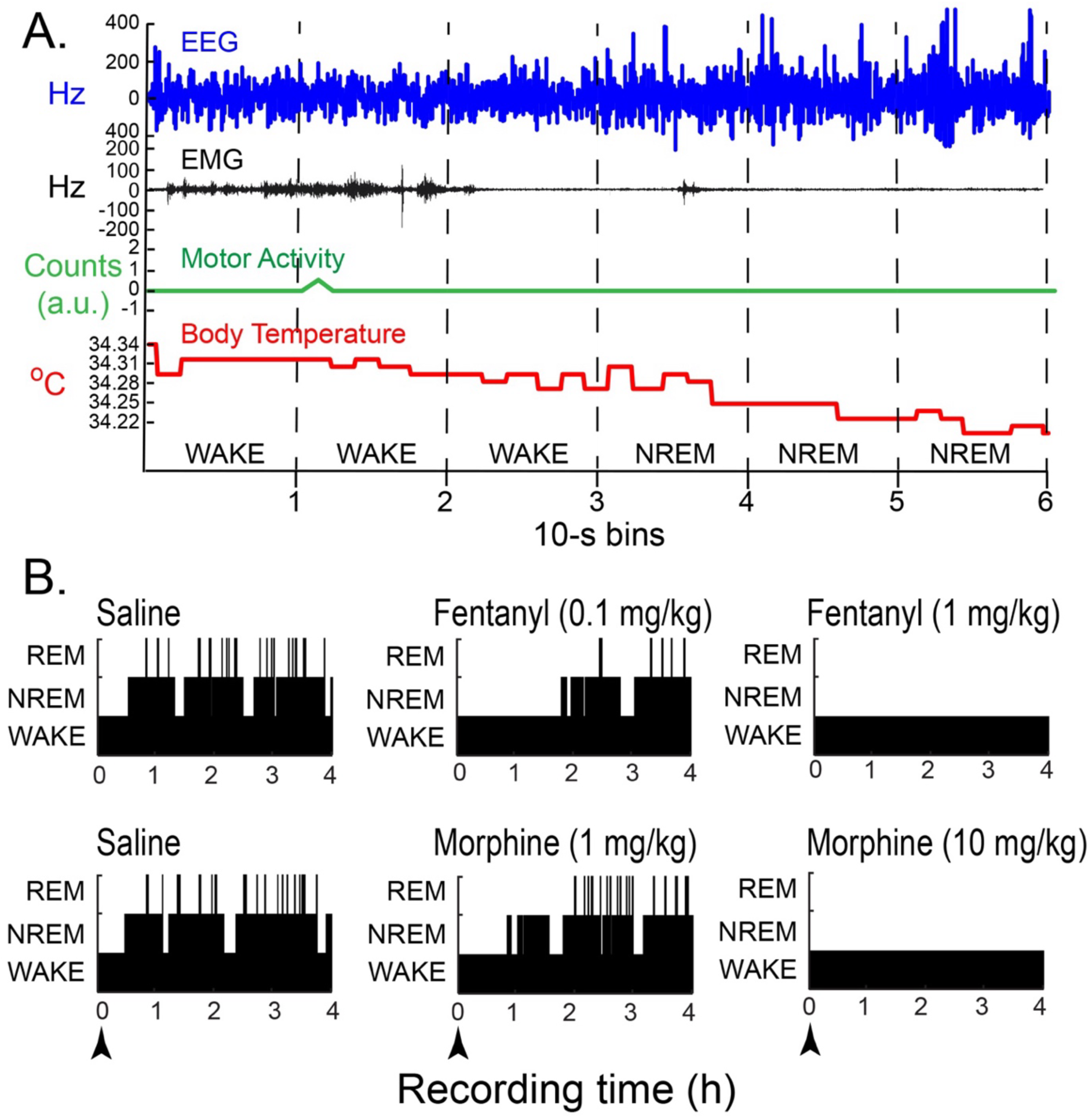
Opioid administration alters neurobehavioral traits and sleep/wake states. (A) Simultaneous recordings of cortical electroencephalogram (EEG), electromyogram (EMG), motor activity (a.u., arbitrary units), and subcutaneous body temperature (°C, degrees Centigrade) from one representative mouse after an injection of saline (vehicle control). Behavioral states were scored in 10-s bins (dashed, vertical lines) during wakefulness (WAKE, bins 1, 2 & 3) and non-rapid eye movement sleep (NREM, bins 4, 5 & 6). (B) These time course plots show the sequence of WAKE (lowest height black bars), NREM sleep (intermediate height black bars), and REM sleep (highest black bars) episodes from one representative mouse during 4 h of recording after the mouse received injections (arrowheads at time 0) of saline (vehicle control), two doses (mg/kg) of fentanyl (top row), and two doses of morphine (bottom row). The lower dose of each opioid decreased both states of sleep compared to its respective control. The highest dose of each opioid eliminated sleep for at least 4 h.

Ponemah software (v6.33, DSI) was used to digitize EEG, EMG, motor activity, and body temperature signals at a sampling frequency of 500 Hz. Bandpass filters of 0.3 to 30 Hz and 10 to 100 Hz were applied to the EEG and EMG signals, respectively. A notch filter of 60 Hz was applied to both signals to reduce electrical noise. The digitized biopotentials were scored offline using NeuroScore™ (v3.3.1, DSI).

Behavioral states of sleep and wakefulness were quantified every 10-s for each 4-h recording (Fig. 1A). The dependent measures were evaluated by two independent scorers, one of whom was blinded to the treatment condition. Agreement between the two scorers was equal to or greater than 90%. Criteria used to score each 10-s bin as wakefulness, NREM sleep, or REM sleep have been previously described (Flint et al., 2010; O’Brien et al., 2021). Briefly, episodes of wakefulness were characterized by a relatively low amplitude, fast frequency EEG signal and a relatively high amplitude EMG signal. Epochs of NREM sleep were distinguished by increased EEG delta activity (0.5 to 4 Hz) and reduced EMG amplitude. REM sleep was identified by increased EEG theta activity (4 to 8 Hz) and a further decrease in EMG amplitude that indicated skeletal muscle atonia. Motor activity and body temperature were analyzed using a MATLAB custom script that averaged activity counts across the 4-h recording period and averaged body temperature during wakefulness, NREM sleep, and REM sleep. A time course plot generated from the sequence of each 10-s epoch (Fig. 1B) was made for every 4-h recording.

### Statistical analyses

Sample sizes were set a priori based on our previous opioid studies using B6 mice (Glovak et al., 2017; Angel et al., 2018; O’Brien et al., 2021). Opioid dose was treated as the independent variable throughout the study. Dependent variables included tail withdrawal latency, percent of recording time spent in states of wakefulness, NREM sleep, and REM sleep, number of episodes and episode duration for each behavioral state, time from injection onset to enter NREM sleep and REM sleep (latency), number of transitions between states, motor activity, and body temperature. These dependent measures were analyzed using Prism (v10.2.3, GraphPad, San Diego, CA) or Statistical Analysis System (SAS V9.4 TS1M7, Cary, NC) for descriptive and inferential statistics. Normality of these data was evaluated using Shapiro-Wilk and Q-Q plots. Effect sizes were calculated for all reported inferential statistics using eta squared (1^2^) and Cohen’s d. All results with a probability (P) value of < 0.05 are reported as statistically significant.

The dose-response effects of fentanyl and morphine Figs. 2A, 3, 5A&D) were analyzed using the non-linear regression equation Y= B + (T-B)/(1+10^(LogED50-X)). Terms in this equation were defined as: “B”, the lower limit of the dependent variable; “T”, the highest limit for the dependent variable response; “X”, the logarithm of the fentanyl or morphine dose; and “Y”, the dependent variable. Nonlinear regression was used to calculate the half-maximal opioid dose (ED_50_) for increasing tail withdrawal latency, wakefulness, and motor activity, and the half-maximal dose for inhibiting (ID_50_) NREM sleep and REM sleep.

**Fig. 2.**
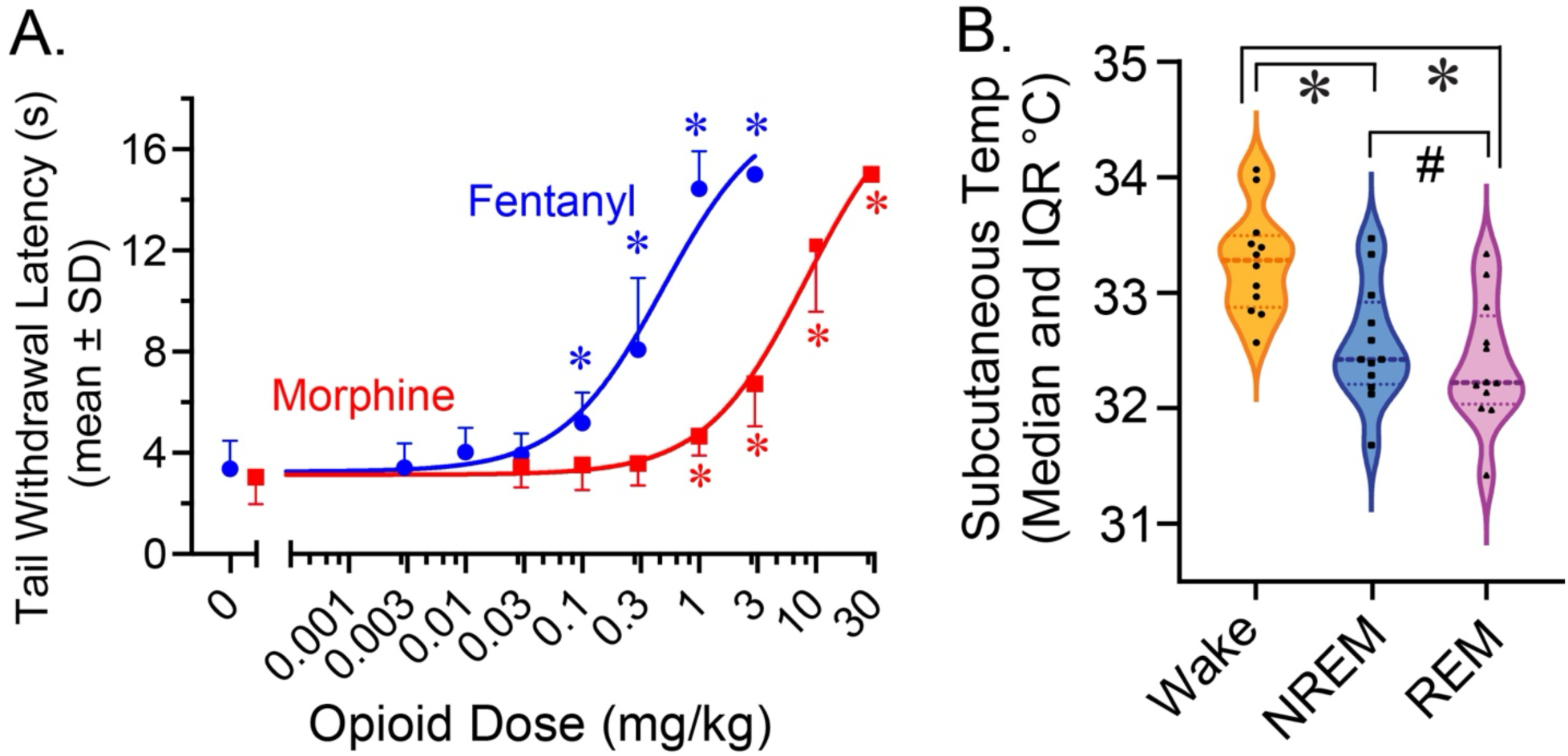
Opioid-induced antinociception and subcutaneous body temperature in adult, male C57BL/6J mice. (A) Tail withdrawal latency was measured in 12 mice, 30 min after administration of saline (0 mg/kg) and seven doses of fentanyl (blue circles) and morphine (red squares). One-way repeated measures ANOVA and Dunnett’s multiple comparisons test identified doses of fentanyl (*P< 0.01) and morphine (*P< 0.02) that caused a significant increase in tail withdrawal latency compared to the respective control (0 mg/kg) for each opioid. (B) Subcutaneous body temperature averaged by sleep/wake state across the 4-h recording period during wakefulness (orange), NREM sleep (blue), and REM sleep (purple) after administration of saline in 12 additional mice. Black filled symbols plot the mean body temperature of each mouse. The dashed, horizontal line within each violin shows median body temperature for all mice. The thin dotted lines mark the 25^th^ and 75^th^ percentile of each distribution. Body temperature data were analyzed by one-way repeated measures ANOVA and Tukey’s multiple comparisons test. Body temperature during NREM sleep and REM sleep was significantly reduced below waking levels (*P < 0.0001). During REM sleep, body temperature was significantly lower than during NREM sleep (^#^P = 0.0052).

**Figure 3.**
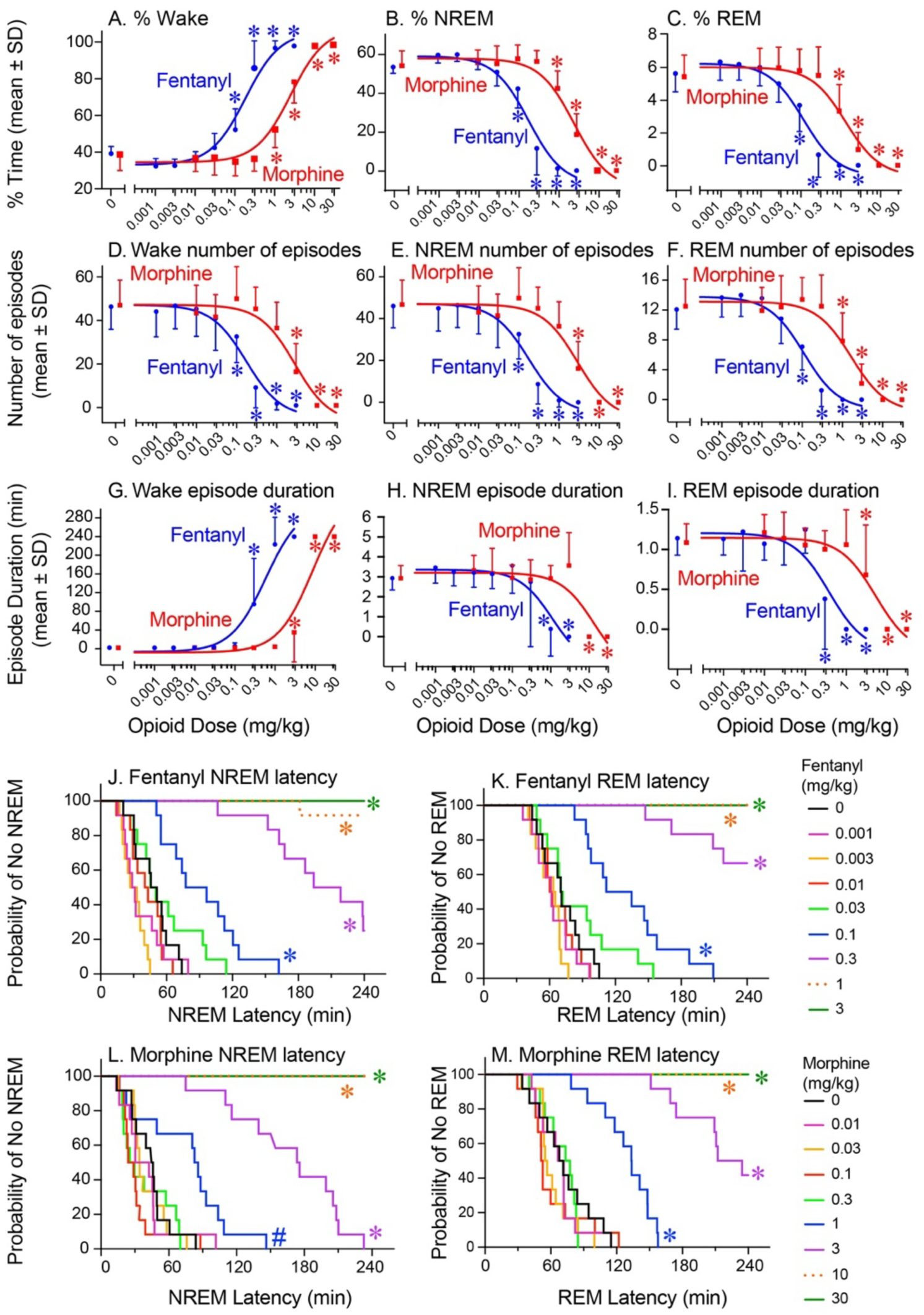
Dose-dependent effects of opioids on sleep and wakefulness. All dependent measures were obtained from 12 mice after administration of saline (0 mg/kg) and eight doses of fentanyl and morphine. (A-I) Dose-response curves show mean ± SD for the 4-h post-injection recoding period. Data were evaluated for statistical significance using one-way repeated measures ANOVA. Dunnett’s multiple comparisons test identified significant differences (*P < 0.05) from saline (vehicle control). (J-M) Survival curves plot the amount of time post-injection (min) until each mouse entered its first NREM or REM sleep episode. There is one curve per dose that represents the average data from all mice. The key at right shows the color code for each dose. Latency data were analyzed by one-way repeated measures ANOVA. Post hoc Dunnett’s multiple comparisons test identified doses of each opioid that caused a significant difference (*P < 0.0001, # P = 0.0046) from saline (0 mg/kg, black curve).

Dose-response data were evaluated for significance using repeated measures, one-way ANOVA and post hoc Dunnett’s multiple comparisons test (Figs. 2A, 3A-M, 5A&D). The effects of opioids on sleep latency were graphed (Figs. 3J-M) using the Kaplan-Meier method for survival analysis (Prism v10.2.3). Repeated measures, two-way ANOVA and post hoc Tukey’s multiple comparisons test evaluated time-by-dose opioid effects on percent state (Fig. 4). Body temperature data were evaluated for significance using repeated measures, one-way ANOVA and post hoc Tukey’s test (Fig 2B).

**Figure 4.**
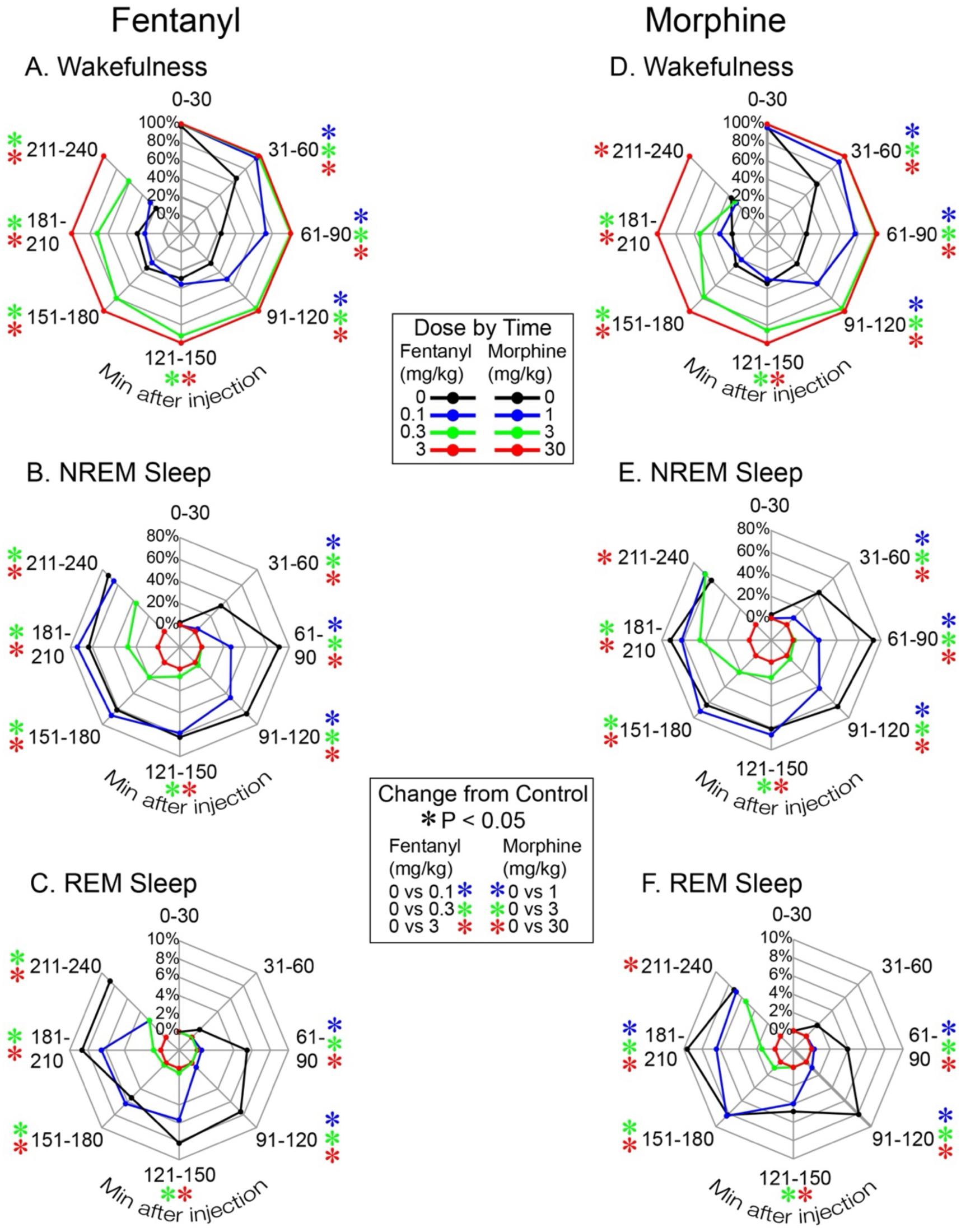
Dose-dependent time course of opioid effects on percent of the 4-h recording period spent in wakefulness, NREM sleep, and REM sleep. Radar plots show the mean amount of time (%) every 30 min during the 4-h recording period after injection of fentanyl (left column) and morphine (right column) that mice (n=12) spent in wakefulness (A & D), NREM sleep (B & E), and REM sleep (C & F). The upper key (Dose by Time) identifies the opioid dose by color. 0 mg/kg indicates saline (vehicle control). The lower key (Change from Control) identifies doses of opioid that caused statistically significant (*P < 0.05) changes from saline for each of the eight 30-min bins. Asterisk color identifies the opioid dose. Data were analyzed by two-way repeated measures ANOVA and post hoc Tukey’s multiple comparison test. These data provide a nuanced view of the relationship between opioid dose and duration of effect.

A within-subject multilevel mediation regression model (Vuorre and Bolger, 2018) was performed using Stata (StataCorp. 2023. *Stata Statistical Software: Release 18*. College Station, TX: StataCorp LLC). These analyses assessed the possible mediation effects of motor activity and body temperature on the relationship between opioid dose and percent of the 4-h recording time spent in wakefulness. For the present study a direct effect indicates how the opioid treatment impacted any one of the dependent variables without influence of a mediator variable. An indirect effect quantifies how the opioid dose altered a dependent variable through a mediator variable. The total effect of fentanyl or morphine is represented by the sum of the direct and indirect effects. A Sobel test was performed to assess the significance of mediation effects for each model.

Statistical assumptions for the mediation analyses were evaluated as follows. Shapiro-Wilk test and Q-Q plots were used to perform normality tests of residuals for all regression models. Outliers and high leverage points were evaluated using the residuals of each model. Heteroskedasticity was evaluated using residual vs fitted plots. Motor activity data underwent a natural log (ln) transformation in order to meet the statistical assumptions of the model. After motor activity was transformed, all statistical assumptions were met.

## Results

### Dose-dependent antinociception

Figure 2A summarizes the dose-response relationships for fentanyl- and morphine-induced antinociception. Tail withdrawal latency data were evaluated by one-way repeated measures ANOVA. There was a significant (P < 0.0001) main effect of fentanyl dose (F_7, 77_ = 190.4) and morphine dose (F_7, 77_ = 154.8) on tail withdrawal latency. The 1^2^ was 0.92 for dose of fentanyl and dose of morphine. Dunnett’s multiple comparisons tests revealed that tail withdrawal latency was significantly increased above control levels by the four highest doses of fentanyl (*P < 0.01), and by the four highest doses of morphine (*P < 0.02). Cohen’s d values for significant doses of both opioids were > 1.

The lowest antinociceptive doses for fentanyl and morphine were 0.1 and 1 mg/kg, respectively. ED_50_ values for fentanyl and morphine were 0.49 and 8.29, respectively, indicating that, as expected, fentanyl was more potent than morphine in providing antinociception. The Fig. 2A results are directly relevant to subsequent figures showing that the lowest dose of fentanyl and morphine that produced antinociception also disrupted sleep. The lowest antinociceptive dose of fentanyl also increased motor activity.

### Sleep-dependent changes in subcutaneous body temperature

Figure 2B plots state-dependent changes in body temperature of B6 mice after injection of saline (vehicle control). These plots confirm that body temperature across the sleep/wake cycle was measured reliably in freely moving mice using the HD-X02 telemeter (Sun et al., 2023). Repeated measures one-way ANOVA indicated that body temperature varied significantly across the sleep wake cycle (F_2, 22_ = 202.8; P < 0.0001; 1^2^= 0.39). Tukey’s multiple comparisons test revealed that body temperature was significantly (*P < 0.0001) decreased below waking levels during NREM sleep and during REM sleep, and that body temperature during REM sleep was significantly lower than body temperature during NREM sleep (^#^P = 0.0052). Cohen’s d values for wakefulness vs NREM sleep and for wakefulness vs REM sleep were >1. Cohen’s d for NREM sleep vs REM sleep was 0.3.

### Fentanyl and morphine reduced time in NREM and REM sleep

Figure 3A shows that the percent of the 4-h recording period spent in wakefulness was progressively increased by increases in opioid dose and was accompanied by concomitant decreases in NREM sleep and REM sleep (Fig. 3B-C). For all 12 mice, sleep was eliminated for at least 4 h by the highest dose of fentanyl (3 mg/kg) and by the two highest doses of morphine (10 and 30 mg/kg). The ED_50_ values for increasing wakefulness were 0.18 for fentanyl and 2.43 for morphine. The ID_50_ values for decreasing NREM sleep were 0.19 for fentanyl and 2.56 for morphine. REM sleep ID_50_ values for fentanyl and morphine were 0.13 and 1.50, respectively. Averaged together, the foregoing ED_50_ and ID_50_ values indicate that fentanyl was approximately 13 times more potent than morphine for increasing wakefulness and decreasing sleep.

The Fig. 3A-C data were analyzed using one-way repeated measures ANOVA to evaluate significant dose effects of fentanyl and morphine. There was a significant (P < 0.0001) treatment main effect for fentanyl on percent of the 4-h recording period spent in wakefulness (Fig. 3A, F8, 88 = 192.3; 1^2^ = 0.9), NREM sleep (Fig. 3B, F8, 88 = 184.1; 1^2^ = 0.9), and REM sleep (Fig. 3C, F8, 88 = 124.0; 1^2^ = 0.9). Dunnett’s multiple comparisons test indicated that doses of fentanyl ≥ 0.1 mg/kg significantly (*P < 0.001) increased percent wakefulness and decreased percent NREM (*P < 0.001) and REM sleep (*P < 0.0001) relative to control (0 mg/kg). Cohen’s d for the significant doses of fentanyl were >1.

There was also a significant (P < 0.0001) treatment main effect of morphine on percent of the 4-h recording period spent in wakefulness (Fig. 3A, F8, 88 = 182.0; 1^2^ = 0.9), NREM sleep (Fig. 3B, F8, 88 = 180.5; 1^2^ = 0.9), and REM sleep (Fig. 3C, F8, 88 = 88.41; 1^2^ = 0.8). Dunnett’s multiple comparisons test revealed that doses of morphine ≥ 1 mg/kg significantly increased percent time in wakefulness (*P < 0.001) and decreased percent time in NREM sleep (*P < 0.001) and REM sleep (*P < 0.0001) relative to control (0 mg/kg). Cohen’s d values for the significant doses of morphine were >1.

### Fentanyl and morphine decreased the number of sleep and wakefulness episodes

Figures 3D-F show that both opioids caused dose-dependent decreases in the number of episodes of wakefulness, NREM sleep, and REM sleep. The ID_50_ values for fentanyl-induced decreases in the number of episodes were 0.17, 0.18, and 0.11 for wakefulness, NREM sleep, and REM sleep, respectively. ID_50_ values for morphine-induced decreases in the number of episodes of wakefulness, NREM sleep, and REM sleep were 2.86, 2.94, and 1.69, respectively. When averaged together, these ID_50_ values indicate that fentanyl was approximately 16 times more potent than morphine for decreasing the number of behavioral state episodes.

One-way repeated measures ANOVA and post hoc Dunnett’s multiple comparisons test were used to identify doses of fentanyl and morphine that significantly changed the number of episodes for each behavioral state. There was a significant (P < 0.0001) treatment main effect for fentanyl on the number of episodes of wakefulness (Fig. 3D, F8, 88 = 80.16; 1^2^ = 0.8), NREM sleep (Fig. 3E, F8, 88 = 82.72; 1^2^ = 0.8), and REM sleep (Fig. 3F, F8, 88 = 97.65; 1^2^ = 0.9). Dunnett’s multiple comparisons test showed that doses of fentanyl ≥ 0.1 mg/kg significantly (*P < 0.001) decreased the number of episodes of all three behavioral states. Cohen’s d for the significant doses of fentanyl were >1.

Significant (P < 0.0001) treatment main effects were found for morphine on the number of episodes of wakefulness (Fig. 3D, F8, 88 = 44.63; 1^2^ = 0.7), NREM sleep (Fig. 3E, F8, 88 = 46.29; 1^2^ = 0.7), and REM sleep (Fig. 3F, F8, 88 = 63.83; 1^2^ = 0.8). Dunnett’s multiple comparisons test revealed that doses of morphine ≥ 3 mg/kg significantly (*P < 0.0001) decreased the number of episodes of wakefulness (Fig. 3D) and NREM sleep (Fig. 3E). The number of REM sleep episodes (Fig. 3F) was significantly (*P < 0.0001) decreased by doses of morphine ≥ 1 mg/kg. Cohen’s d for the significant doses of morphine were >1.

### Fentanyl and morphine increased the duration of waking episodes

The dose-dependent decreases in the number of episodes of wakefulness, NREM sleep, and REM sleep described above were accompanied by concomitant increases in the duration of wakefulness episodes (Fig. 3G), and corresponding decreases in NREM and REM sleep episode duration (Fig. 3H-I). The ED_50_ values for opioid-induced increases in duration of wakefulness episodes were 0.59 for fentanyl and 8.70 for morphine, respectively (Fig. 3G). ID_50_ values for opioid-induced decreases in duration of NREM sleep episodes were 1.11 for fentanyl and 13.6 for morphine (Fig. 3H). ID_50_ values for opioid-induced decreases in REM sleep episode duration were 0.38 for fentanyl and 5.25 for morphine (Fig. 3I). These ED_50_ and ID_50_ values, when averaged together, indicate that fentanyl was approximately 14 times more potent than morphine for effects on duration of episodes for all three behavioral states.

One-way repeated measures ANOVA showed a treatment main effect for fentanyl on duration of episodes for wakefulness (Fig. 3G, F8, 88 = 85.28, P < 0.0001; 1^2^ = 0.9), NREM sleep (Fig. 3H, F8, 88 = 17.67, P < 0.0001; 1^2^ = 0.5), and REM sleep (Fig. 3I, F8, 88 = 37.38, P < 0.0001; 1^2^ = 0.7). Dunnett’s multiple comparisons test indicated that doses of fentanyl ≥ 0.3 mg/kg significantly (*P < 0.0001) increased episode duration of wakefulness and decreased REM sleep episode duration. NREM sleep episode duration was significantly (*P < 0.0001) decreased by the two highest doses of fentanyl, 1 and 3 mg/kg. Cohen’s d for the significant doses of fentanyl were >1.

There was a treatment main effect of morphine on duration of wakefulness episodes (Fig. 3G, F8, 88 = 287.1, P < 0.0001; 1^2^ = 0.9), NREM sleep (Fig. 3H, F8, 88 =34.18, P < 0.0001; 1^2^ = 0.7), and REM sleep (Fig. 3I, F8, 88 = 30.88, P < 0.0001; 1^2^ = 0.7). Dunnett’s multiple comparisons test indicated that the 3, 10, and 30 mg/kg doses of morphine significantly increased the duration of waking episodes (*P < 0.02) and decreased the duration of REM sleep episodes (*P < 0.02). The 10 and 30 mg/kg doses of morphine decreased duration of NREM sleep episodes (*P < 0.0001).

Cohen’s d values for the significant doses of morphine were >0.8, with the exception of duration of wakefulness episodes at the 3 mg/kg dose, which had a Cohen’s d = 0.7. In summary, both opioids disrupted sleep architecture by significantly increasing the duration of wakefulness episodes (Fig. 3G), thereby decreasing the number of wakefulness episodes (Fig. 3D) and decreasing the number and duration of sleep episodes (Fig. 3E, F, H, and I).

### Sleep latency was increased by antinociceptive doses of fentanyl and morphine

Survival curves (Fig. 3J-M) plot the dose-dependent increases in latency to onset of NREM sleep and REM sleep caused by fentanyl and morphine. The probability of NREM sleep (Fig. 3J) and REM sleep (Fig. 3K) decreased with increasing doses of fentanyl. The highest dose of fentanyl (3 mg/kg) increased NREM sleep latency to greater than 240 min for all 12 mice (Figure 3J ). The two highest doses of morphine (10 and 30 mg/kg) increased latency to NREM sleep (Fig. 3L) and to REM sleep (Fig. 3M) latency to greater than 240 min for all 12 mice.

Nonlinear regression analyses were applied to the dose-response data for latency to onset of NREM and REM sleep. ED_50_ values for the opioid-induced increases in NREM sleep latency were 0.18 for fentanyl and 2.95 for morphine. ED_50_ values for REM sleep latencies were 0.14 for fentanyl and 1.62 for morphine. These results indicate that fentanyl was approximately 14 times more potent than morphine for inhibiting the onset of both NREM and REM sleep.

One-way repeated measures ANOVA showed a treatment main effect for fentanyl on latency to onset of NREM sleep (Fig. 3J, F8, 88 = 189.9, P < 0.0001; 1^2^ = 0.9) and REM sleep (Fig. 3K, F8, 88 = 181.6, P < 0.0001; 1^2^ = 0.9). Dunnett’s multiple comparisons test indicated that doses of fentanyl ≥ 0.1 mg/kg significantly (*P < 0.0001) increased latency to onset of NREM and REM sleep. Cohen’s d for the significant doses of fentanyl were >1.

One-way repeated measures ANOVA indicated a treatment main effect for morphine on latency to onset of NREM sleep (Fig. 3L, F8, 88 =191.3, P < 0.0001; 1^2^ = 0.9) and REM sleep (Fig. 3M, F8, 88 = 203.4, P < 0.0001; 1^2^ = 0.9). Dunnett’s multiple comparisons test indicated that doses of morphine ≥ 1 mg/kg significantly increased NREM sleep latency (*P < 0.01) and REM sleep latency (*P < 0.001). Cohen’s d values for the significant doses of morphine were >1.

### Fentanyl and morphine changed the number of transitions between states of sleep and wakefulness

As illustrated by Fig. 1B, mouse sleep is characterized by four types of state transitions: wakefulness to NREM sleep, NREM sleep to REM sleep, REM sleep to wakefulness, and NREM sleep to wakefulness. The numbers of these four types of state transitions are reported in Table 1 and were analyzed using repeated measures one-way ANOVA with multiple comparisons to saline (0 mg/kg) provided by Dunnett’s (*P < 0.01). Table 1A summarizes the results for fentanyl. One-way ANOVA showed a significant (P < 0.0001) main effect of fentanyl dose on number of transitions from wakefulness to NREM sleep (F_2, 22_ = 72.4; 1^2^ = 0.7), NREM sleep to REM sleep (F_2, 22_ = 56.1; 1^2^ = 0.8), REM sleep to wakefulness (F_2, 22_ = 49.9; 1^2^ = 0.7), and NREM sleep to wakefulness (F_2, 22_ = 55.62; 1^2^ = 0.6). For morphine (Table 1B) there was a significant (P < 0.001) main effect of dose on number of transitions from wakefulness to NREM sleep (F_2, 22_ = 23.9; 1^2^ = 0.5), NREM sleep to REM sleep (F_2, 22_ = 54.1; 1^2^ = 0.6), REM sleep to wakefulness (F_2, 22_ = 54.53; 1^2^ = 0.6), and NREM sleep to wakefulness (F_2, 22_ = 12.0; 1^2^ = 0.4).

**TABLE 1.**
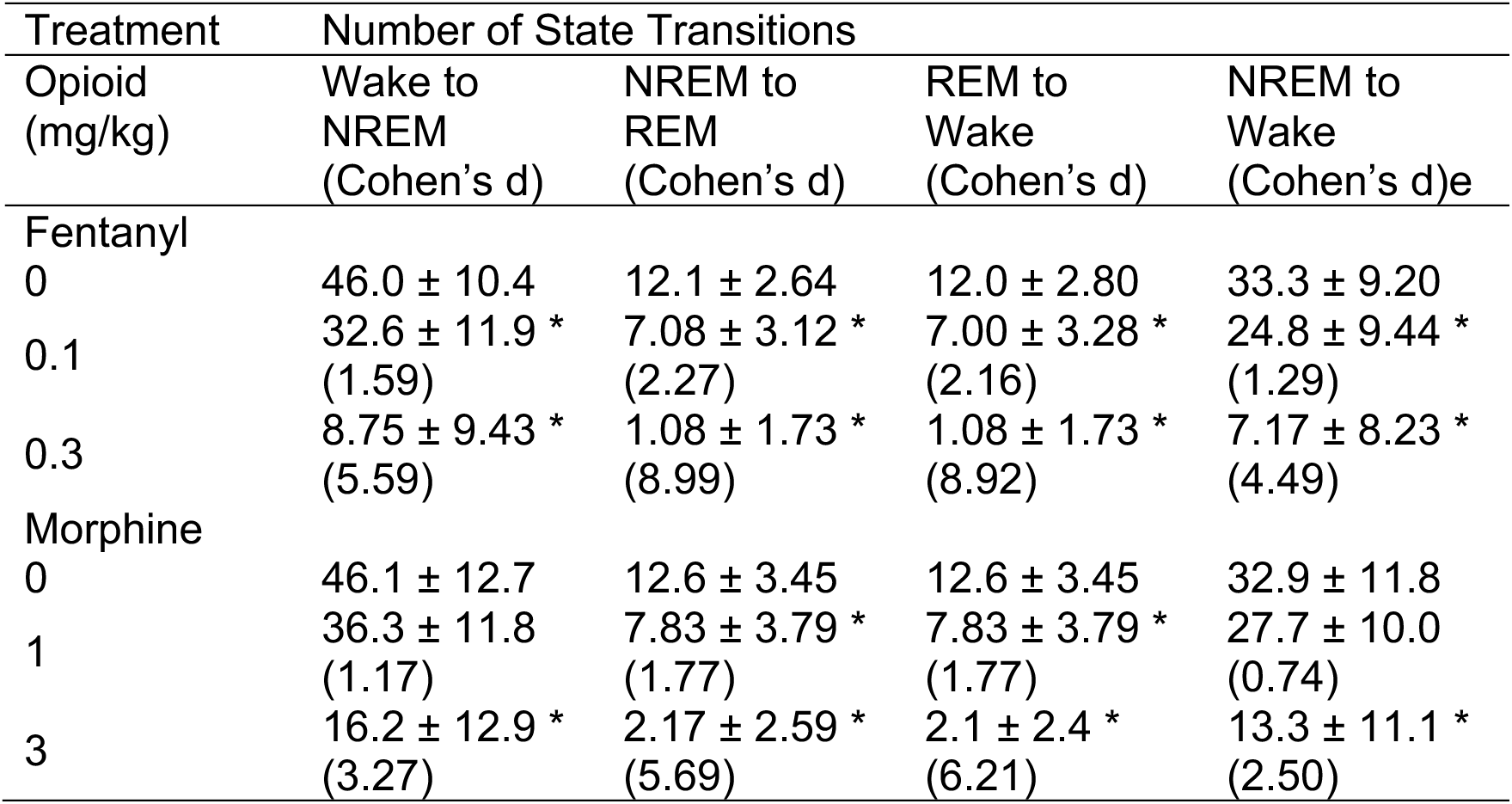
Mean ± SD number of transitions between states of wakefulness (Wake), rapid eye movement sleep (REM), and non-REM sleep (NREM). Four types of state transitions were quantified for 4 h after mice (n=12) received saline (vehicle control, 0 mg/kg), and the lowest antinociceptive doses and the next highest doses of fentanyl (0.1, 0.3 mg/kg) and morphine (0, 1, 3 mg/kg). Data were analyzed using repeated measures one-way ANOVA (see Results section for F values, degrees of freedom, P values, and effect sizes for the ANOVAs). Multiple comparisons to saline (0 mg/kg) were calculated using Dunnett’s (*P < 0.01).

| Treatment | Number of State Transitions |  |  |  |
| --- | --- | --- | --- | --- |
| Opioid (mg/kg) | Wake to NREM<br>(Cohen's d) | NREM to REM<br>(Cohen's d) | REM to Wake<br>(Cohen's d) | NREM to Wake<br>(Cohen's d)e |
| Fentanyl |  |  |  |  |
| 0 | 46.0 ± 10.4 | 12.1 ± 2.64 | 12.0 ± 2.80 | 33.3 ± 9.20 |
| 0.1 | 32.6 ± 11.9 *<br>(1.59) | 7.08 ± 3.12 *<br>(2.27) | 7.00 ± 3.28 *<br>(2.16) | 24.8 ± 9.44 *<br>(1.29) |
| 0.3 | 8.75 ± 9.43 *<br>(5.59) | 1.08 ± 1.73 *<br>(8.99) | 1.08 ± 1.73 *<br>(8.92) | 7.17 ± 8.23 *<br>(4.49) |
| Morphine |  |  |  |  |
| 0 | 46.1 ± 12.7 | 12.6 ± 3.45 | 12.6 ± 3.45 | 32.9 ± 11.8 |
| 1 | 36.3 ± 11.8<br>(1.17) | 7.83 ± 3.79 *<br>(1.77) | 7.83 ± 3.79 *<br>(1.77) | 27.7 ± 10.0<br>(0.74) |
| 3 | 16.2 ± 12.9 *<br>(3.27) | 2.17 ± 2.59 *<br>(5.69) | 2.1 ± 2.4 *<br>(6.21) | 13.3 ± 11.1 *<br>(2.50) |

### Time course of dose-dependent opioid-induced sleep disruption

The Fig. 4 radar plots show the time course of sleep and wakefulness after injection of saline (black lines, 0 mg/kg), the lowest effective antinociceptive doses of fentanyl and morphine (blue lines, 0.1 and 1 mg/kg, respectively) and the next highest doses of fentanyl and morphine (green lines, 0.3 and 3 mg/kg, respectively). Red lines illustrate changes in behavioral state caused by the highest tested doses of fentanyl (3 mg/kg) and morphine (30 mg/kg). Each 4-h recording period is illustrated as eight 30-min bins.

The dose by time effects of fentanyl are illustrated by Figs. 4A-C and were analyzed by two-way repeated measures ANOVA. For the percent of the 4-h recording time spent in wakefulness (Fig. 4A), there was a significant (P < 0.0001) time main effect (F_7, 77_ = 44.14; 1^2^ = 0.2), dose main effect (F_3, 33_ = 98.57; 1^2^ = 0.4), and time by dose interaction (F_21, 231_ = 11.16; 1^2^ = 0.1). For percent of time spent in NREM sleep (Fig. 4B) there was a significant (P < 0.0001) time main effect (F_7, 77_ = 40.1; 1^2^ = 0.2), dose main effect (F_3, 33_ = 97.8; 1^2^ = 0.4), and time by dose interaction (F_21, 231_ = 10.3; 1^2^ = 0.1). For the percent of the total recording time spent in REM sleep (Fig. 4C) there was a significant (P < 0.0001) time main effect (F_7, 77_ = 20.85; 1^2^ = 0.1), dose main effect (F_3, 33_ = 66.63; 1^2^ = 0.3), and time by dose interaction (F_21, 231_ = 5.62; 1^2^ = 0.1).

The dose by time effects of morphine are illustrated by Figs. 4D-F and were analyzed by two-way repeated measures ANOVA. For percent of time spent in wakefulness (Fig. 4D) there was a significant (P < 0.0001) time main effect (F_7, 77_ = 48.21; 1^2^ = 0.2), dose main effect (F_3, 33_ = 139.2; 1^2^ = 0.4), and time by dose interaction (F_21, 231_ = 13.16; 1^2^ = 0.2). For NREM sleep (Fig. 4E) there was a significant (P < 0.0001) time main effect (F_7,77_ = 46.86; 1^2^ = 0.2), dose main effect (F_3,33_ = 147.1; 1^2^ = 0.4), and time by dose interaction (F_21, 231_ = 12.63; 1^2^ = 0.2). For REM sleep (Fig. 4F) there was a significant (P < 0.0001) time main effect (F_7, 77_ = 28.24; 1^2^ = 0.2), dose main effect (F_3, 33_ = 61.39; 1^2^ = 0.2), and time by dose interaction (F_21, 231_ = 6.41; 1^2^ = 0.2).

### Mediation analyses revealed that opioid-induced increases in wakefulness were partially mediated by increased motor activity but not by changes in subcutaneous body temperature

Figure 5A plots the increase in motor activity caused by increasing doses of fentanyl and morphine relative to control (0 mg/kg). The ED_50_ values for the effects of fentanyl and morphine on motor activity were 0.10 and 5.88 mg/kg, respectively. One-way repeated measures ANOVA showed a significant (P < 0.0001) main effect of fentanyl (F_8, 88_ = 34.7; 1^2^ = 0.6) and morphine (F_8, 88_ = 59.24; 1^2^ = 0.7) on motor activity during wakefulness. Dunnett’s multiple comparisons test revealed that doses of fentanyl ≥ 0.1 mg/kg and doses of morphine ≥ 3 mg/kg significantly (*P <0.001) increased motor activity. Cohen’s d values for the significant doses of fentanyl and morphine were >1.

**Figure 5.**
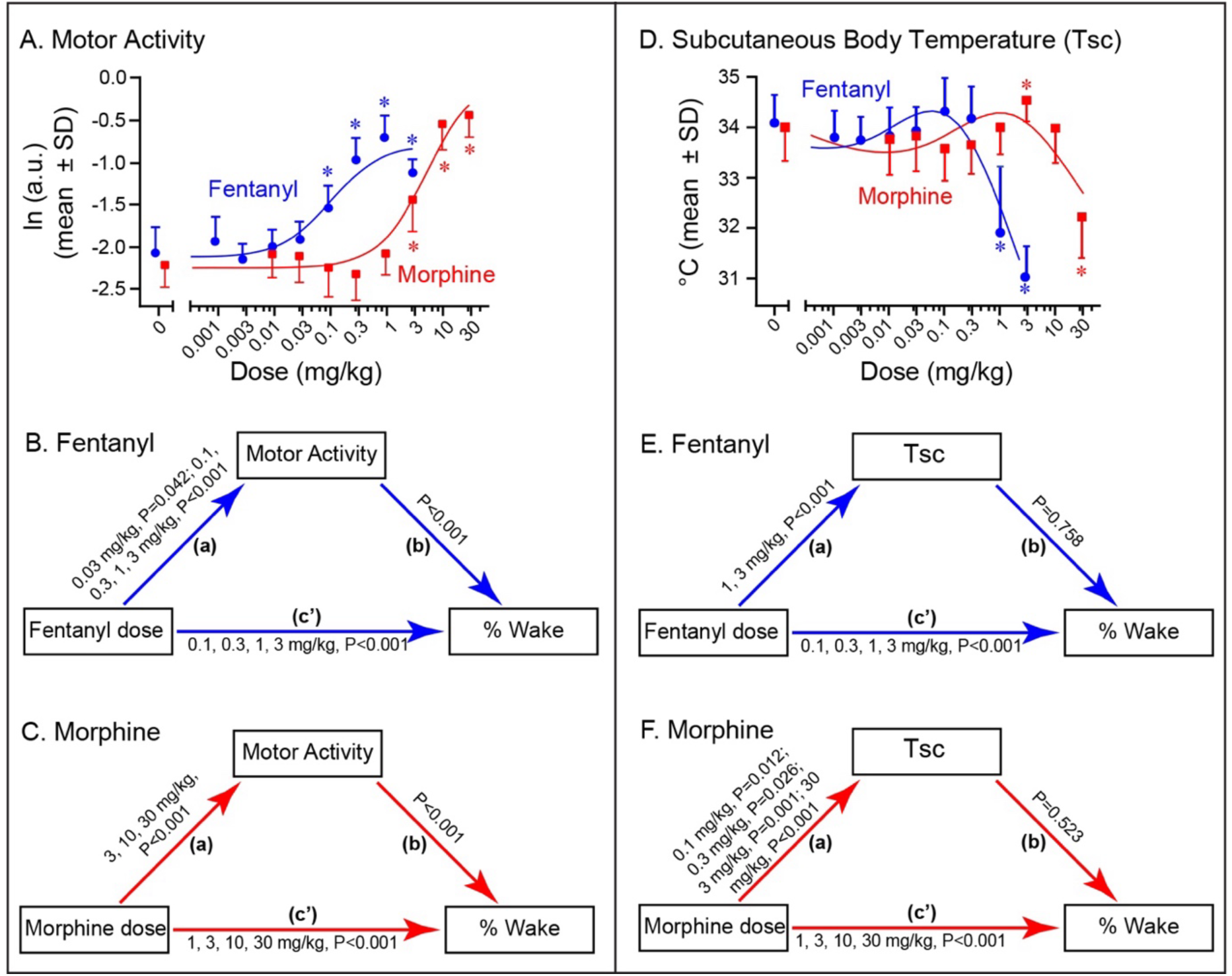
Dose-response curves and mediation analyses for opioid effects on motor activity and subcutaneous body temperature. The two upper graphs (A and D) illustrate results from repeated measures ANOVA and Dunnett’s multiple comparisons test identifying doses of fentanyl (blue circles) and morphine (red squares) that caused a significant (**\***P< 0.05) change from control (0 mg/kg) for (A) motor activity plotted as the natural logarithm (ln) of activity counts in arbitrary units (a.u.) and (D) subcutaneous body temperature plotted as degrees centigrade (°C). The four lower plots (B-C and E-F) are path diagrams for mediation analyses illustrating the relationships between the direct, indirect, and total effects of opioid dose, motor activity and body temperature on percent of time spent in wakefulness. For the path analysis of motor activity (B-C) the blue and red arrows indicate causal paths for (**a**) the effect of opioid dose on motor activity, (**b**) the effect of motor activity on %Wake, and (**c’**) the effect of opioid dose on %Wake with motor activity considered. Significant opioid doses (mg/kg) with their respective P values are listed above each of the three arrows (**a, b, c’**) for effects of fentanyl dose (B) and morphine dose (C). The path analysis of body temperature (E-F) illustrates the model used to estimate the mediation effect of opioid-induced changes in body temperature on opioid-induced increases in wakefulness. For the E-F plots, the arrows indicate causal paths (**a**) for the effect of opioid dose on body temperature, (**b**) the effect of body temperature on %Wake, and (**c’**) the effect of opioid dose on %Wake with body temperature considered. Significant opioid doses (mg/kg) with their respective P values are listed above each of the three arrows (**a, b, c’**) for effects of fentanyl dose (E) and morphine dose (F). Mediation model diagrams were modified from Vuorre and Bolder, 2018.

Figure 5B and 5C summarize the results of within-subject multi-level mediation analysis used to assess whether motor activity affected the relationship between opioid dose and percent of time spent awake. Opioid dose (mg/kg) was the independent variable hypothesized to be causal, percent of time spent in wakefulness (% Wake) was the main dependent measure, and motor activity was hypothesized to have mediated the effect of opioid dose on percent of the 4-h recording time spent in wakefulness.

The upward pointing blue arrow **(a)** at left of Fig. 5B indicates that fentanyl doses ≥ 0.03 mg/kg significantly increased motor activity. The downward pointing blue arrow **(b)** at right of Fig. 5B indicates a significant effect of motor activity on % Wake. The horizontal blue arrow **(c’)** of Fig. 5B illustrates the findings that when taking motor activity into account, fentanyl dose was a significant predictor of % Wake for doses ≥ 0.1 mg/kg. For every 1% increase in motor activity, the model predicted a 0.5% increase in % Wake. A Sobel test determined significant mediation (P < 0.001) with 15.9% of the variance in % Wake mediated by motor activity.

Figure 5C shows the results of mediation analysis for effects of morphine on motor activity. The upward pointing red arrow at left (**a**) revealed that doses ≥ 3 mg/kg significantly increased motor activity. The downward pointing red arrow **(b)** at right identifies the significant effect of motor activity on % Wake. The horizontal red arrow **(c’)** shows that when motor activity was considered, morphine doses ≥ 1 mg/kg were significant predictors of % Wake. For every 1% increase in motor activity, the model predicted a 0.8% increase in % Wake. A Sobel test determined significant mediation (P < 0.0001) with 21.7% of the variance in % Wake mediated by motor activity.

Figure 5D plots the dose-dependent changes in subcutaneous body temperature during wakefulness caused by fentanyl and morphine. The ID_50_ value for the effect of fentanyl on body temperature was 2.78 mg/kg. The body temperature data for morphine did not fit the nonlinear regression model. The spline function in Prism was used to fit both curves without a model.

Figure 5D summarizes the findings that one-way repeated measures ANOVA showed a significant (P < 0.0001) main effect of fentanyl (F_8, 88_ = 59.3; 1^2^ = 0.7) and morphine (F_8, 88_ = 25.7; 1^2^ = 0.5) on body temperature during wakefulness. Dunnett’s multiple comparisons test revealed that the two highest doses of fentanyl (1 and 3 mg/kg) significantly (*P < 0.001) decreased body temperature. Higher doses of morphine had a biphasic effect on body temperature. Dunnett’s multiple comparisons test showed that morphine at 3 mg/kg significantly increased body temperature (*P = 0.02), and at 30 mg/kg morphine significantly decreased body temperature (*P < 0.0001). Cohen’s d values for the significant doses of fentanyl and morphine were >0.9.

Within-subject multi-level mediation regression analysis also was used to determine whether body temperature influenced the relationship between opioid dose and percent of the 4-h recording time spent awake. As described above, opioid dose (mg/kg) was the independent variable hypothesized to be causal, percent of time spent in wakefulness (% Wake) was the main dependent measure, and body temperature was proposed to have mediated the effect of opioid dose on % Wake. In contrast to motor activity, changes in body temperature did not mediate the effect of fentanyl dose or morphine dose on wakefulness. Figure 5E shows that fentanyl at doses of 1 and 3 mg/kg significantly altered body temperature (upward pointing blue arrow (**a**) at left), but there was no significant effect of body temperature on % Wake (downward pointing blue arrow (**b**) at right). The horizontal blue arrow (**c’**) shows that when body temperature was included as a covariate in the analysis, fentanyl at doses ≥ 0.1 mg/kg significantly increased % Wake.

In Fig. 5F, the upward pointing red arrow (**a**) shows the morphine doses that significantly altered body temperature, and the downward pointing red arrow (**b**) reports a nonsignificant effect of body temperature on % Wake. The horizontal red arrow (**c’**) shows that when body temperature was considered in the analyses, morphine at doses ≥ 1 mg/kg significantly increased % Wake. In summary, mediation analyses revealed that the fentanyl- and morphine-induced increases in wakefulness were partially mediated (15.9% and 21.7%, respectively) by motor activity (Fig. 5C), but not by opioid-induced changes in body temperature (Fig. 5F).

## Discussion

Three novel findings shaped this discussion. First, antinociceptive doses of acutely administered fentanyl and morphine disrupted sleep architecture by increasing wakefulness and reducing NREM and REM sleep. Second, fentanyl and morphine caused dissociations among sleep/wake states, body temperature, and motor activity by lowering body temperature while increasing wakefulness and motor activity. Third, sleep disruption was partially mediated by the opioid-induced increase in motor activity but not by opioid-induced changes in body temperature.

### Antinociceptive doses of fentanyl and morphine disrupt sleep architecture

Sleep architecture refers to the predictable sequence of sleep states across the sleep period. Sleep fragmentation is a disruption of sleep architecture defined by the intrusion of short episodes of wakefulness during the sleep period (Ramesh et al., 2009; Greenwald et al., 2021; Li et al., 2022). Sleep architecture in mice (Fig. 1B, Fig. 3, Table 1) and humans (Wang and Teichtahl, 2007) is assessed by quantifying the amount of time spent in each state of sleep, latency to sleep onset, number and duration of NREM and REM episodes, and number and type of transitions between states.

Opioids prescribed for clinical management of pain commonly produce undesired effects, including disrupting sleep architecture (Eacret et al., 2020; Greenwald et al., 2021; Lydic et al., 2022; Akinnusi et al., 2024). Even a single dose of morphine significantly disrupts sleep architecture in healthy adults (Shaw et al., 2005; Dimsdale et al., 2007). Sleep-disrupting effects of opioids exacerbate pain (Roehrs et al., 2006; Muncey et al., 2010; Dalanon et al., 2021). Sleep disruption contributes to poorer cardiometabolic outcomes (Altman et al., 2012) and longer recovery periods after brain injury (Fleming et al., 2020). Sleep disruption may also contribute to relapse of opioid use disorder (Fathi et al., 2020; James et al., 2020; Tripathi et al., 2020b; Eckert and Yaggi, 2022; Huhn and Finan, 2022; Lydic et al., 2022). The foregoing unwanted effects of opioids are produced by complex interactions among multiple neuronal networks involved in regulating sleep, pain (Haack et al., 2020), and opioid craving during withdrawal (Greenwald et al., 2021; Eckert and Yaggi, 2022).

The present dose-response data revealed the novel finding that even the lowest effective, antinociceptive doses of fentanyl and morphine caused significant increases in wakefulness and decreases in NREM sleep and REM sleep (Fig. 3). Antinociceptive doses of fentanyl and morphine also significantly decreased the number of state transitions (Table 1). Thus, in B6 mice, diminishing nociception by administering opioids cannot be achieved without disrupting sleep for at least 4 h (Fig. 4).

Benzodiazepines and antidepressants have different effects on NREM sleep than on REM sleep (Hambrecht-Wiedbusch et al., 2010; Roehrs and Roth, 2010; Dhuna and Malkani, 2020). In contrast, doses of fentanyl and morphine that increased wakefulness decreased NREM and REM sleep. The findings that antinociceptive doses of both opioids increased the duration of wakefulness episodes, decreased the number of episodes of all states, and decreased transitions between behavioral states show that in B6 mice opioids significantly disrupt every aspect of sleep architecture for up to 4 h (Fig. 4).

### Endogenous changes in mouse body temperature

Mammalian body temperature exhibits a circadian rhythm, and sleep onset in humans occurs during the maximal rate of decline in body temperature (Czeisler et al., 1980; Campbell and Broughton, 1994). Body temperature also fluctuates during sleep (Glotzbach and Heller, 1976) and is lower during NREM sleep than during wakefulness in birds and mammals (Heller, 1988). Figure 2B shows the expected decrease in body temperature during NREM sleep compared with wakefulness under control conditions. Body temperature also decreased below waking and NREM sleep temperatures during REM sleep, consistent with the finding that all mammals studied to date do not regulate their body temperature during REM sleep (Glotzbach and Heller, 1976; Heller, 1988; Jhaveri et al., 2007; Tan and Knight, 2018). Regarding effects of ambient temperature on mouse body temperature, the B6 mice in the present study were housed at ambient temperatures that did not significantly change NREM or REM sleep during the light period (Jhaveri et al., 2007).

### Fentanyl and morphine decreased body temperature while increasing motor activity and wakefulness

Body temperature did not show a classic sigmoidal, agonist dose response to morphine (Fig. 5D). Both opioids at higher doses significantly decreased body temperature despite causing significant increases in locomotor activity (Fig. 5A). Motor activity is thermogenic (Abreu-Vieira et al., 2015) and normally precedes an increase in mouse body temperature (Lateef et al., 2014). Morphine had a biphasic effect on body temperature, whereas fentanyl decreased body temperature (Fig. 5B). These dose-dependent effects on body temperature are similar to rectal temperature changes reported for NMRI mice in response to similar doses of morphine and fentanyl (Baker and Meert, 2002). The results show that fentanyl dissociated the normal physiological relationship of motor activity to body temperature. Fentanyl has been shown to dissociate the normal relationship between motor activity and breathing in B6 mice by increasing locomotor activity while decreasing minute ventilation (Furdui et al., 2023).

Dose-dependent increases in motor activity caused by fentanyl and morphine in B6 mice have been documented (Santos et al., 2022; Olson et al., 2023; Varshneya et al., 2023). Assays of motor activity typically involve administering a drug before placing a mouse in a novel environment and quantifying the number of times an infrared beam is broken and/or distance traveled (Fowler, 2010). A novel aspect of the present study is that states of sleep and wakefulness, motor activity, and body temperature were measured simultaneously while mice remained in their home cages. This approach avoided the increase in exploratory behavior that results in sleep disruption when mice are placed in a novel environment (Tang et al., 2005).

The lowest effective doses of fentanyl and morphine that increased motor activity (Fig. 5A) were also the lowest effective doses for disrupting sleep (Figs. 3&4). These novel findings are consistent with the interpretation that opioids disrupt interactions among neural circuits regulating body temperature, motor activity, and states of sleep and wakefulness. Mediation analysis was then used to determine whether opioid-induced sleep disruption was caused by increased motor activity and/or changes in body temperature.

### Opioid-induced sleep disruption was partially mediated by motor activity

Mediation analysis is a statistical method used to quantify the extent to which intervening variables (mediators) affect an outcome (Vuorre and Bolger, 2018). This approach was used to determine whether opioid-induced changes in motor activity and/or body temperature contributed to the dose-dependent increase in wakefulness caused by opioids. The results showed that the increases in motor activity induced by specific doses of fentanyl and morphine mediated a portion of the opioid-induced increase in wakefulness (Fig. 5). These data also revealed that although both opioids significantly changed body temperature, those changes did not mediate a significant portion of the opioid-induced increases in wakefulness.

### Limitations, future directions, and conclusions

The data were limited to male B6 mice and focused on fentanyl as a widely used drug of abuse, and morphine as the gold standard for mu opioid receptor mediated effects. These results encourage future studies of female B6 mice to characterize sex-specific and dose-dependent effects of fentanyl, morphine, and opioids used to treat opioid use disorder, such as methadone and buprenorphine (Angel et al., 2018; Glovak et al., 2022). Single-dose studies show that buprenorphine disrupts sleep in male B6 mice (O’Brien et al., 2021) male Sprague Dawley rats (Gauthier et al., 2011), and men taking buprenorphine to treat opioid use disorder (Tripathi et al., 2020a). Single-dose opioids are a common feature of post-operative pain management. The 4-h duration of the present recordings (Fig. 4) is consistent with typical surgery times for human (Crespin et al., 2022) and veterinary (Shaver and Hofmeister, 2018) medicine.

Future studies of chronic opioid administration are needed to determine whether tolerance develops to the sleep-disruptive effects of chronic opioid administration in mice. In patients taking methadone or buprenorphine for medication assisted therapy, REM sleep is suppressed, and daytime sleepiness has been reported (Greenwald et al., 2021). The present results encourage studies of dose-related effects of chronic opioid administration on multiple, simultaneously collected physiological variables to provide insights into how chronically used opioids affect sleep and interacting physiological control systems.

In conclusion, the present data quantify the effects of fentanyl and morphine on sleep states and neurobehavioral traits in mice. Similarities to human data include sleep disruption, hyperthermia, and hypothermia. An important species-specific difference in mice is the opioid-induced increase of motor activity. This study demonstrated that mediation analyses can differentiate the effects of opioids on wakefulness in the presence of other intervening factors. Mediation analyses that incorporate the complexity of opioids causing direct and indirect effects have provided insights relevant to opioid-induced respiratory depression (Orr et al., 2024) and clinical management of pain (Whibley et al., 2019). The present study is a novel application of mediation analysis to quantify opioid-induced changes in the neurobehavioral function of B6 mice. The results also illustrate the power of a systems pharmacology approach (Bai and Abenethy, 2013) for unmasking opioid-induced, dose-dependent changes in clinically relevant sleep/wake states, nociception, motor activity, and body temperature.

## Data Availability

All the data supporting the findings of this paper are available within the paper.

## Abbreviations

EEG: electroencephalogram /electroencephalographic
B6: C57BL/6J
Tsc: subcutaneous body temperature
EMG: electromyogram
REM: rapid eye movement
NREM: non-rapid eye movement
ED_50:_: half-maximal dose for increasing a response
ID_50:_: half-maximal dose for decreasing a response
ln: natural log

## Acknowledgments

Preliminary data quantifying the dose-response effects of fentanyl and morphine on states of sleep and wakefulness, motor activity, and body temperature in B6 mice have been presented (Zebadúa Unzaga et al., 2021; Zebadúa Unzaga et al., 2022; Cooper et al., 2023).

## References

Abreu-Vieira G, Xiao C, Gavrilova O and Reitman ML (2015) Integration of body temperature into the analysis of energy expenditure in the mouse. Mol Metab 4:461–470.

Akinnusi M, Martinson A and El-Solh AA (2024) Treatment of insomnia associated with alcohol and opioid use: a narrative review. Sleep Biol Rhythms 22:1–17.

Altman NG, Izci-Balserak B, Schopfer E, Jackson N, Rattanaumpawan P, Gehrman PR, Patel NP and Grandner MA (2012) Sleep duration versus sleep insufficiency as predictors of cardiometabolic health outcomes. Sleep Med Clin 13:1261–1270.

Anderson EJ and Jeter JP (2023) Dermatological implications of sleep deprivation in the US military. Cutis 111:146–149.

Angel C, Glovak ZT, Alami W, Mihalko S, Price J, Jiang Y, Baghdoyan HA and Lydic R (2018) Buprenorphine depresses respiratory variabillity in obese mice with altered leptin signaling. Anesthesiology 128:984–991.

Atwood ME (2022) Effects of sleep deficiency on risk, course, and treatment of psychopathology. Clin Chest Med 43:305–318.

Bai JPF and Abenethy DR (2013) Systems pharmacology to predict drug toxicity: Integration across levels of biological organization. Annu Rev Pharmacol Toxicol 53:451–473.

Baker AK and Meert TF (2002) Functional effects of systemically administered agonists and antagonists of μ, δ, and κ opioid receptor subtypes on body temperature in mice. J Pharmacol Exp Ther 302:1253–1264.

Baldo BA (2025) Opioid-induced respiratory depression: clinical aspects and pathophysiology of the respiratory network effects. Am J Physiol Lung Cell Mol Physiol 328:L267–L289.

Brower KJ and Perron BE (2010) Sleep disturbance as a universal risk factor for relapse in addictions to psychoactive substances. Med Hypotheses 74:928–933.

Campbell SS and Broughton RJ (1994) Rapid decline in body temperature before sleep: fluffing the physiological pillow? Chronobiol Int 11:126–131.

Cipriano A, Apseloff G, Kapil RP, He E, Shet M and Harris SC (2025) Time course of reversal of fentanyl-induced respiratory depression in healthy subjects by intramuscular almefene and intramuscular and intranasal naloxone. J Clin Pharm 65:206–216.

Collins CL, England K, Conrad SW, Day H, Tilahun E, Pan GD and Meyer T (2025) Patient-reported duration of opioid analgesic use after discharge from surgical procedures or other types of acute pain: a scoping review. Pain Med 26:1–12.

Cooper NW, Zebadúa Unzaga D, Olson CE, Garrett GJ and Baghdoyan HA (2023) Fentanyl and morphine decrease sleep duration without causing sleep fragmentation in C57BL/6J mice. Soc Neurosci Abstracts Program No PSTR55706.

Crespin DJ, Ruder T, Mulcahy AW and Mehrota A (2022) Variation in estimated surgical procedure times across patient characteristics and surgeon specialty. JAMA Surg 157:e220099.

Czeisler CA, Weitzman ED, Moore-Ede MC, Zimmerman JC and Knauer RS (1980) Human sleep: Its duration and organization depend on its circadian phase. Science 210:1264–1267.

Dahlhamer J, Lucas J, Zelaya C, Nahin R, Mackey S, DeBar L, Kerns R, Von Korff M, Porter L and Helmick C (2018) Prevalence of chronic pain and high-impact chronic pain among adults - United States, 2016. Morb Mortal Wkly Rep 67:1001–1006.

Dalanon J, Chikahisa S, Shiuchi T, Shimizu N, Chavan P, Suzuki Y, Okura K, Séi H and Matsuka Y (2021) Pain sensitivity increases with sleep disturbance under predictable chronic mild stress in mice. Sci Rep 11:14231.

Dhuna NA and Malkani RG (2020) Antidepressants and their impact on sleep. Curr Sleep Medicine Rep 6:216–225.

Dimsdale JE, Norman D, DeJardin D and Wallace MS (2007) The effect of opioids on sleep architecture. J Clin Sleep Med 3:33—36.

Djonlagic I, Mariani S, Fitzpatrick AL, Klei VMvd, Johnson DA, Wood AC, Seeman T, Nguyen HT, Prerau MJ, Luchsinger JA, Dzierzewski JM, Rapp SR, Tranah GJ, Yaffe K, Burdick KE, Stone KL, Redline S and Purcell SM (2021) Macro and micro sleep architecture and cognitive performance in older adults. Nat Human Behav 5:123–125.

Dowell D, Ragan K, Jones C, Baldwin G and Chou R (2022) CDC clinical practice guideline for prescribing opioids for pain — United States. Morb Mortal Wkly Rep 71:1–95.

Duerksen JM, Ramjiawan M, Witt J, Fitzpatrick-Wong S, Tappia PS, Ramjiawan B, Mansouri B, Sareen J and Knight E (2025) The addictive process of opioids: current and novel interventions in opioid use disorder. Can J Physiol Pharmacol 103:111–122.

Eacret D, Veasey SC and Blendy JA (2020) Bidirectional relationship between opioids and disrupted sleep: Putative mechanisms. Mol Pharmacol 98:445–453.

Eckert DJ and Yaggi HK (2022) Opioid use disorder, sleep deficiency, and ventilatory control: Bidirectional mechanisms and therapeutic targets. Am J Respir Crit Care Med 206:937–949.

Fathi HR, Yoonessi A, Khatibi A, Rezaeitalab F and Rezaei-Ardani A (2020) Crosstalk between sleep disturbance and opioid use disorder: a narrative review. Addict Health 12:140–158.

Finan PH, Goodin BR and Smith MT (2013) The association of sleep and pain: an apdate and a path forward. J Pain 14:1539–1552.

Fleming MK, Smejka T, Henderson Slater D, Van Gils V, Garratt E, Yilmaz Kara E and Johansen-Berg H (2020) Sleep disruption after brain injury is associated with worse motor outcomes and slower functional recovery. Neurorehabil Neural Repair 34:661–671.

Fletcher D and Martinez V (2014) Opioid-induced hyperalgesia in patients after surgery: a systematic review and a meta-analysis. Br J Anaesth 112:991–1004.

Flint RR, Chang T, Lydic R and Baghdoyan HA (2010) GABA-A receptors in the pontine reticular formation of C57BL/6J mouse modulate neurochemical, electrographic, and behavioral phenotypes of wakefulness. J Neurosci 30:12301–12309.

Fowler SC (2010) Motor activity and stereotypy, in *Encyclopedia of Psychopharmacology* (Stolerman IP ed) pp 799-805, Springer, Berlin.

Furdui A, Scarpellini CdS and Montandon G (2023) Fentanyl-induced respiratory depression and locomotor hyperactivity are mediated by mu-opioid receptors. eNeuro 10:1–18.

Gamble MC, Chuan B, Gallego-Martin T, Shelton MA, Puig S, O’Donnell CP and Logan RW (2022) A role for the circadian transcription factor NPAS2 in the progressive loss of non-rapid eye movement sleep and increased arousal during fentanyl withdrawal in male mice. Psychopharmacology 239:3185–3200.

Gårdmark M, Höglund AU and Hammarlund-Udenaes M (1998) Aspects on tail-flick, hot-plate and electrical stimulation tests for morphine antinociception. Pharmacol Toxicol 83:252–258.

Gauthier EA, Guzick SE, Brummett CM, Baghdoyan HA and Lydic R (2011) Buprenorphine disrupts sleep and decreases adenosine concentrations in sleep-regulating brain regions of Sprague Dawley rat. Anesthesiology 115:743–753.

Glare P, Mahendran M and Weiss AD (2025) Update on chronic pain management. Intern Med J 55:200–206.

Glotzbach SF and Heller HC (1976) Central nervous system regulation of body temperature during sleep. Science 194:537–539.

Glovak Z, Mihalko S, Baghdoyan HA and Lydic R (2017) Leptin status alters buprenorphine-induced antinociception in obese mice with dysfunctional leptin receptors. Neurosci Lett 660:29–33.

Glovak ZT, Angel C, O’Brien CB, Baghdoyan HA and Lydic R (2022) Buprenorphine differentially alters breathing among four congenic mouse lines as a function of dose, sex, and leptin status. Respir Physiol Neurobiol 297:1–9.

Greenwald MK, Moses TE and Roehrs TA (2021) At the intersection of sleep deficiency and opioid use: mechanisms and therapeutic opportunities. Transl Res 234:58–73.

Haack M, Simpson N, Sethna N, Kaur S and Mullington J (2020) Sleep deficiency and chronic pain: potential underlying mechanisms and clinical implications. Neuropsychopharmacology 45:205–216.

Hambrecht-Wiedbusch VS, Gauthier EA, Baghdoyan HA and Lydic R (2010) Benzodiazepine receptor agonists cause drug-specific and state-specific alterations in EEG power and acetylcholine release in rat pontine reticular formation. Sleep 33:909–918.

Haouzi P, McCann M and Tubbs N (2021) Respiratory effects of low and high doses of fentanyl in control and beta-arrestin 2-deficient mice. J Neurophysiol 125:1396–1407.

Heller HC (1988) Sleep and hypometabolism. Can J Zool 66:61–69.

Huhn AS and Finan PH (2022) Sleep disturbance as a therapeutic target to improve opioid use disorder treatment. Exp Clin Psychopharmacol 30:1024–1035.

James MH, Fragale JE, Aurora RN, Cooperman NA, Langleben DD and Aston-Jones G (2020) Repurposing the dual orexin receptor antagonist suvorexant for the treatment of opioid use disorder: why sleep on this any longer? Neuropsychopharmacology 45:717–791.

Jhaveri KA, Trammell RA and Toth LA (2007) Effect of environmental temperature on sleep, locomotor activity, core body temperature and immune responses of C57BL/6J mice. Brain Behav Immun 21:975–987.

Knutson K (2015) Sleep and pain: summary of the 2015 Sleep in America Poll. Sleep Health 1:85.

Lateef DM, Abreu-Vieira G, Xiao C and Reitman ML (2014) Regulation of body temperature and brown adipose tissue thermogenesis by bombesin receptor subtype-3. Am J Physiol Endocrinol Metab 306:E681–E687.

Li S-B, Damonte VM, Chen C, Wang GX, Kebschull JM, Yamaguchi H, Bian W-J, Purmann C, Pattni R, Urban AE, Mourrain P, Kauer JA, Scherrer G and Lecea Ld (2022) Hyperexcitable arousal circuits drive sleep instability during aging. Science 375:1–14.

Little KM and Kosten TA (2023) Focus on fentanyl in females: Sex and gender differences in the physiological and behavioral effects of fentanyl. Front Neruoendocrinol 71:1–11.

López-Muciño LA, García-García F, Cueto-Escobedo J, Acosta-Hernández M, Venebra-Muñoz A and Rodríguez-Alba JC (2022) Sleep loss and addiction. Neurosci Biobehav Rev 141:1–11.

Lydic R, Hillman DR, Jiang Y, Baghdoyan HA and O’Brien CO (2022) Opioid actions on sleep and breathing, in Principles and Practice of Sleep Medicine (Kryger MH, Roth T and Goldstein CA eds) pp 510–518, Elsevier, Philidelphia.

Marchette RCN, Vendruscolo LF and Koob GF (2025) The dynorphin/-opioid receptor rystem at the interface of hyperalgesia/hyperkatifeia and addiction. Curr Add Rep 12:1–14.

McLain JM, Alami WH, Glovak ZT, Cooley CR, Burke SJ, Collier JJ, Baghdoyan HA, Karlstad MD and Lydic R (2018) Sleep fragmentation delays wound healing in a mouse model of type 2 diabetes. Sleep 41:1–9.

Mickley GA, Mulvihill MA and Postler MA (1990) Brain mu and delta opioid receptors mediate different locomotor hyperactivity responses of the C57BL/6J mouse. Psychopharmacology 101:332–337.

Morrell M, Finn L, Kim H, Peppard P, Badr MS and Young T (2000) Sleep fragmentation, awake blood pressure, and sleep-disordered breathing in a population-based study. Am J Respir Crit Care Med 162:2091–2096.

Muncey AR, Saulles AR, Koch LG, Britton SL, Baghdoyan HA and Lydic R (2010) Disrupted sleep and delayed recovery from chronic peripheral neuropathy are distinct phenotypes in a rat model of metabolic syndrome. Anesthesiology 113:1176–1185.

O’Brien CO, Locklear C, Glovak ZT, Zebadúa Unzaga D, Baghdoyan HA and Lydic R (2021) Opioids cause dissociated states of consciousness in C57BL/6J mice. J Neurophysiol 126:1265–1275.

Olson KM, Hillhouse TM, Burgess GE, West JL, Hallahan JE, Dripps IJ, Ladetto AG, Rice KC, Jutkiewicz EM and Traynor JR (2023) Delta opioid receptor-mediated antidepressant-like effects of diprenorphine in mice. J Pharmacol Exp Ther 384:343–352.

Orr JE, Malhotra A, Gruenberg E, Marin T, Sands SA, Alex RM, Owens RL and Schmickl CN (2024) Pathogenesis of sleep-disordered breathing in the setting of opioid use: a multiple mediation analysis using physiology. Sleep 47:1–9.

Paxinos G and Franklin K (2019) The Mouse Brain in Stereotaxic Coordinates. Academic Press, San Diego.

Percie du Sert N, Hurst V, Ahluwalia A, Alam S, Avey MT, Baker M and al. e (2020) The ARRIVE guidelines 2.0: Updated guidelines for reporting animal research. PLOS Biology 18:e3000410.

Piantadosi S (1997) *Clinical trials: A methodologic perspective*. Wiley, New York.

Ramesh V, Kaushal N and Gozal D (2009) Sleep fragmentation modifies EEG delta power during slow wave sleep in socially isolated and paired mice. Sleep Science 2:64–75.

Ramirez JM, Burgraff NJ, Wei AD, Baertsch NA, Varga AG, Baghdoyan HA, Lydic R, Morris KF, Bolser DC and Levitt ES (2021) Neuronal mechanisms underlying opioid-induced respiratory depression: our current understanding. J Neurophysiol 125:1899–1919.

Roehrs T, Hyde M, Blaisdell B, Greenwald M and Roth T (2006) Sleep loss and REM sleep loss are hyperalgesic. Sleep 29:145–151.

Roehrs T and Roth T (2010) Drug-related sleep stage changes: functional significance and clinical relevance. Sleep Med Clin 5:559–570.

Roehrs T, Sibai M and Roth T (2021) Sleep and alertness disturbance and substance use disorders: A bi-directional relation. Pharmacol Biochem Behav 203:1–8.

Rosow CE, Miller JM, Pelikan EW and Cochin J (1980) Opiates and thermoregulation in mice. I. Agonists. J Pharmacol Exp Ther 213:273–283.

Santos EJ, Banks ML and Negus SS (2022) Role of efficacy as a determinant of locomotor activation by mu opioid receptor ligands in female and male mice. J Pharmacol Exp Ther 382:44–53.

Schildhaus N, Trink E, Polson C, Detolla L, Tyler B, Jallo G, Tok S and Guarnieri M (2014) Thermal latency studies in opiate-treated mice. J Pharm Bioallied Sci 6:43–47.

Sharma R, Parikh M, Chischolm A, Kempuraj D and Thakkar M (2024) Dopamine D2 receptors in the accumbal core region mediates the effects of fentanyl on sleep-wakefulness. Neuroscience 560:11–19.

Shaver SL and Hofmeister EH (2018) Factors affecting the duration of anesthesia and surgery of canine and feline gonadectomies performed by veterinary students in a year-long preclinical surgery laboratory. Vet Surg 48:352–359.

Shaw IR, Lavigne G, Mayer P and Choinière M (2005) Acute intravenous administration of morphine perturbs sleep architecture in healthy pain-free young adults: a preliminary study. Sleep 28:677–682.

Sun Y, Tisdale RK, Yamashita A and Kilduff TS (2023) Peripheral vs. core body temperature as hypocretin/orexin neurons degenerate: Exercise mitigates increased heat loss. Peptides 164:1–10.

Tan CL and Knight ZA (2018) Regulation of body temperature by the nervous system. Neuron 98:31–48.

Tang X, Xiao J, Parris BS, Fang J and Sanford LD (2005) Differential effects of two types of environmental novelty on activity and sleep in BALB/cJ and C57BL/6J mice. Physiol Behav 85:419–429.

Thomas RJ (2024) A matter of fragmentation. Sleep 47:1–4.

Tripathi R, Dhawan A, Rao R, Mishra AK, Jain R and Sinha S (2020a) Assessment of subjective sleep problems in men with pioid dependence maintained on buprenorphine. J Addict Med 14:132–138.

Tripathi R, Rao R, Dhawan A, Jain R and Sinha S (2020b) Opioids and sleep - a review of literature. Sleep Med Clin 67:269–275.

Vanini G, Nemanis K, Baghdoyan HA and Lydic R (2014) GABAergic transmission in rat pontine reticular formation regulates the induction phase of anesthesia and modulates hyperalgesia caused by sleep deprivation. Eur J Neurosci 40:2264–2273.

Varshneya NB, Walentiny DM, Moisa LT, Walker TD, Akinfiresoye LR and Beardsley PM (2019) Opioid-like antinociceptive and locomotor effects of emerging fentanylrelated substances. Neuropharmacology 151:171–179.

Varshneya NB, Walentiny DM, Stevens DL, Walker TD, Akinfiresoye LR and Beardsley PM (2023) Structurally diverse fentanyl analogs yield differential locomotor activities in mice. Pharmacol Biochem Behav 222:1–7.

Volkow ND and Dye LR (2025) Groundbreaking research from NIDA addressing the challenges of the opioid epidemic. J Med Tox 21:69–77.

Vuorre M and Bolger N (2018) Within-subject mediation analysis for experimental data in cognitive psychology and neuroscience. Behav Res Methods 50:2125–2143.

Wang D and Teichtahl H (2007) Opioids, sleep architecture and sleep-disordered breathing. Sleep Med Rev 11:35–46.

Whibley D, AlKandari N, Kristensen K, Barnish M, Rzewuska M, Druce KL and Tang NKY (2019) Sleep and pain A systematic review of studies of mediation. Clin J Pain 35:544–558.

Yamamoto DJ, Woo C-W, Wager TD and Regner MF (2015) Influence ofdorsolateral prefrontalcortex and ventralstriatum on risk avoidance in addiction:vA mediation analysis. Drug Alcohol Depend 149:10–17.

Yamazaki EM, Antler CA, Lasek CR and Goel N (2021) Residual, differential neurobehavioral deficits linger after multiple recovery nights following chronic sleep restriction or acute total sleep deprivation. Sleep 44:zsaa224.

Zebadúa Unzaga D, Bustamante C, Thibert M and Baghdoyan HA (2021) Fentanyl and morphine cause dose-dependent sleep disruption in C57BL/6J (B6) mice. FASEB J 35:1.

Zebadúa Unzaga D, Zhu X, Price J, Lydic R and Baghdoyan HA (2022) Fentanyl-induced sleep disruption in C57BL/6J (B6) mice is not secondary to increased motor activity or hypothermia. FASEB J 36:R2168.

